# Neuron-glia Integrity: Functional Assessment, Molecular Underpinnings, and Implication for Higher Brain Functions

**DOI:** 10.1101/2020.03.23.003681

**Authors:** Haiyan Zeng, Xiaolei Zhang, Wenqiang Wang, Zhiwei Shen, Zhuozhi Dai, Zhijia Yu, Shuqin Xu, Gen Yan, Qingjun Huang, Renhua Wu, Xi Chen, Haiyun Xu

## Abstract

We developed a theory of neuron-glia integrity to underline the fact that neurons and glia cells work together in the central nervous system. Here we substantiated this theory and exemplified the implication of intact neuron-glia integrity for higher brain functions. An animal model of maternal separation with early weaning (MSEW) was applied to neonatal rats to mimic early life neglect and abuse in humans. Behavioral performance of rats was evaluated at adulthood, followed by functional assessments of neuron-glia integrity in living rats, and the demonstration of molecular underpinnings of impaired neuron-glia integrity in MSEW rats. MSEW rats showed higher levels of anxiety and explorative activity, higher glutamate level, but lower GABA level in PFC and hippocampus. MSEW procedure down-regulated protein levels of GLT-1 and ATP-α, but up-regulated GAD65 and GS, while had no effects on GLAST and PAG. Moreover, it reduced the fractional anisotropy values in various brain regions, in addition to increasing NAA levels. Concurrently, MSEW led to hypomyelination in PFC as evidenced by relevant cellular and molecular changes.

## Introduction

Brain cells are classified into neurons and glial cells including astrocytes, microglia, and oligodendrocytes (OLs). Neurons have been thought of as principal cells in the brain because they receive and transmit chemical and electrical signals. Neurons connect each other thus constitute neuronal chains known as neural networks through which electrical signals propagate in the central nervous system (CNS). In this respect, glial cells play an information processing role. For example, astrocytes contact with synapses between neurons and regulate synaptic transmission as the third part of the tripartite synapse (Allen and Barres, 2009; Oberheim et al., 2009). OLs in the brain wrap neuronal axons and form the myelin sheath in the brain. The compacted myelin sheath provides high electrical resistance and low capacitance that is essential for saltatory impulse propagation (Foster et al., 2019). These knowledge supports a notion that astrocyte and neuron function as an integrative entity thereby signal transmission within CNS is adjusted (Murai and Pasquale, 2011; Schousboe et al., 2014) and that axon-myelin co-works as an integrative entity that ensures partly processed information can be properly synchronized at post-synaptic sites which is critical for information processing associated with perception, thought and action (Miller, 2002; Xiao et al., 2016). Based on these advances in neuron-glia communication research, the neuron-glia integrity theory was developed recently to highlight the importance of intact neuron-glia integrity for higher brain functions and mental health (Xu et al., 2016).

According to the neuron-glia integrity theory, the glutamate-glutamine-GABA cycle is the underpinning of neuron-astrocyte entity. Any changes in glutamate (Glu), glutamine (Gln), and/or GABA in a brain region are indicative of an imbalanced Glu-Gln-GABA cycle or impaired neuron-astrocyte entity there. N-acetyl-aspartate (NAA) is considered the neurochemical correlate of axon-myelin entity because it is involved in myelination and axon-glial signaling, in addition to a role in osmoregulation (Moffett et al., 2007; Xu et al., 2016). All these brain metabolites (Glu, Gln, GABA, and NAA), along with the others, can be measured and quantified in living subjects by means of non-invasive neuroimaging methods including proton magnetic resonance spectroscopy (^1^H-MRS) (Maddock and Buonocore, 2012) and chemical exchange saturation transfer (CEST) (Cai et al., 2012). As such, the status of neuron-glia entities can be evaluated and may be related to higher brain functions and brain disorders, including mental disorders (Xu et al., 2016). However, no such a study was published so far that was driven by the neuron-glia integrity theory.

Early-life stresses have been shown to influence brain development and associated with cognitive impairment and affective disorders (Fone and Porkess, 2008). For example, social interactions are indispensable for normal brain development of rats; without such interactions, rats showed changes in behavior (Wilkinson et al., 1994), neurochemistry (Toua et al., 2010; Trabace et al., 2012), and neuroanatomy (Vanderschuren and Trezza, 2014). Indeed, post-weaning social isolation in rats has been employed as an animal model for research relevant to some of mental disorders including anxiety (Regenass et al., 2018), depression (Wang et al., 2017), and schizophrenia (Moller et al., 2011; Witten et al., 2014). Similarly, maternal separation with early weaning (MSEW) paradigm has been used to mimic early life neglect and abuse in humans (George et al., 2010). The latter is believed to influence brain development and consequently bring forth a predisposition toward mental and behavioral disorders (Strüber et al., 2014). Taken together, these previous studies encouraged us to employ an animal model of early life adversity in rats for the demonstration of the neuron-glia integrity theory. We for the first time measured the integrity of neuron-glia entities in brain of living animals by non-invasive neuroimaging techniques including ^1^H-MRS, CEST of glutamate (GluCEST), and diffusion tensor imaging (DTI), demonstrated the molecular underpinnings of impaired neuron-glia integrity in MSEW rats, and linked impaired neuron-glia integrity to abnormalities in higher brain functions of the rats.

## Results

### MSEW increases explorative activity and anxiety levels of rats

After the MSEW procedure (see materials and methods for details), rats continued to live under group housing as those in Control group. On postnatal days (PDs) 60-62, rats in the MSEW and Control groups were subjected to open field test to measure their locomotor activity, exploratory behavior, and anxiety level. It was found: 1) MSEW rats walked a longer distance in whole arena (Fig. 1A) of the open field compared to Control group (6685.50 ±874.12 cm vs 5331.59 ±723. 86 cm, p<0.001); 2) MSEW rats moved at a higher speed (Fig. 1B) compared to Control group (11.24 ± 1.41 cm/s vs 9.04 ± 1.17 cm/s, p<0.001); 3) MSEW rats walked a shorter distance on the central zone (Fig. 1C) compared to Control group (380.64 ± 94.07 cm vs 462.46 ± 64.07 cm, p<0.01); and 4) MSEW rats spent less time on the central zone (Fig. 1D) compared to Control group (21.36 ± 6.69. s vs 27.31 ± 3.93 s, p<0.01). These data indicate the presence of higher levels of explorative activity and anxiety in the MSEW rats relative to Control group.

**Fig.1.**
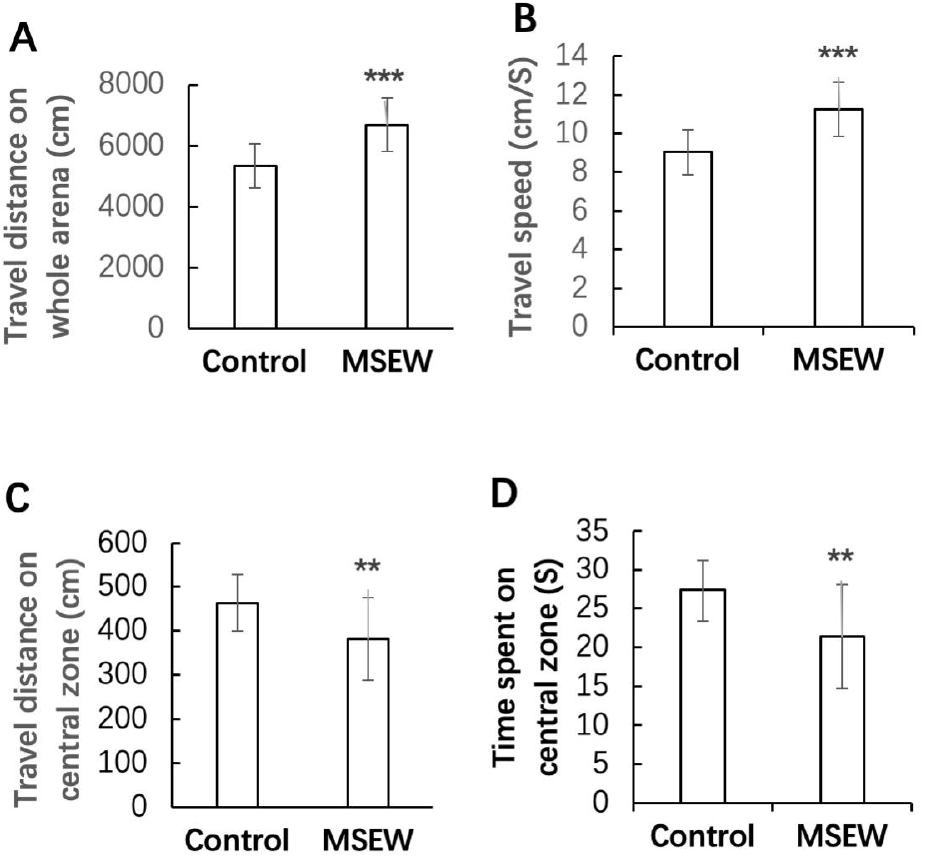
MSEW increased exploratory activity and anxiety level of rats. (**A**) MSEW rats moved a longer distance on whole arena of the open field compared to Control group. (**B**) MSEW rats moved at a higher velocity on the open field compared to Control group. (**C**) MSEW rats moved a shorter distance on the central zone of the open field compared to Control group. (**D**) MSEW rats spent less time on the central zone of the open field compared to Control group. Data are expressed as mean ± SD. N = 15/group. Comparisons were made between Control and MSEW groups. **p<0.01; ***p<0.001.

To examine possible nutritional effects of MSEW procedure on rats, the body weight of rats was weighed on PDs 7, 14, 21, and 30, respectively. MSEW rats were same as those in Control group in terms of body weight measured at all the time points (data not shown), indicating the paradigm did not cause malnutrition in rat pups.

### MSEW elevates global GluCEST contrast in the brain and increases Glu levels in prefrontal cortex and hippocampus of rats

GluCEST imaging has been used to map spatial distribution of Glu in the brain with a good spatial resolution in previous studies (Bagga et al., 2016; Cai et al., 2012; Pépin et al., 2016). As such, it was used in this study to examine possible effects of MSEW on Glu distribution of rats. Before GluCEST imaging, CEST phantoms were prepared and scanned to optimize CEST and MRI procedures. The results showed that the GluCEST contrast was linearly proportional to Glu concentration in the physiological range at pH = 7.0 (Fig. S1), but the other metabolites (NAA, 10 mM; creatine, 6 mM; Gln, 2 mM; Asp, 2 mM; and GABA, 2 mM) contributed negligible CEST effects to GluCEST signal (Fig. S2).

For GluCEST imaging, three horizontal T2-weighted anatomical slice (thickness = 2 mm) images were acquired. Of them, the second one that crosses through the dorsal hippocampus and prefrontal cortex (PFC) was selected as the target slice (Fig. S3A). The hippocampus and PFC are two brain regions essential for higher brain functions and compromised in some of mental disorders including anxiety, depression and schizophrenia (Hiser and et al., 2018; Liu et al., 2017; Penades et al., 2019). The mean B_0_ shift (Fig. S3B, □ 0.5 ppm) and the B_1_/B_1ref_ ratio (Fig. S3C, about 0.5) measured on this slice showed the rather good homogeneity of B_0_ and B_1_ fields. Relative to the GluCEST map of Control group (Fig. S3D), the MSEW rats (Fig. S3E) exhibited GluCEST contrast increase across the whole slice, reflecting a global increase in Glu concentration. The Z-spectra and MTRasym (asymmetrical magnetization transfer ratio) curves also indicated higher GluCEST contrast in the MSEW rats compared to Control group (Fig. S3F).

Following the global GluCEST scanning, regional analysis of GluCEST contrast was done. The regions of interest (ROIs) were PFC (Fig. 2A & B) and dorsal hippocampus (Fig. 2D & E) which were manually drawn based on the T2-weighted reference image of each rat. The corresponding Z-spectra and MTRasym curves of these two ROIs (Fig. 2C & F, respectively) were obtained. The GluCEST maps showed higher GluCEST contrasts in both PFC and hippocampus of MSEW rats (Fig. 2B & E, respectively) compared to those in the Control group (Fig. 2A & D, respectively). This observation was confirmed by Z-spectra and MTRasym curves (Fig. 2C & F). The summarized mean GluCEST contrasts in PFC and hippocampus showed higher GluCEST contrasts in both PFC and hippocampus of MSEW group compared to the Control group (Fig. 2G), suggesting that MSEW increased Glu levels in the two brain regions.

**Fig.2.**
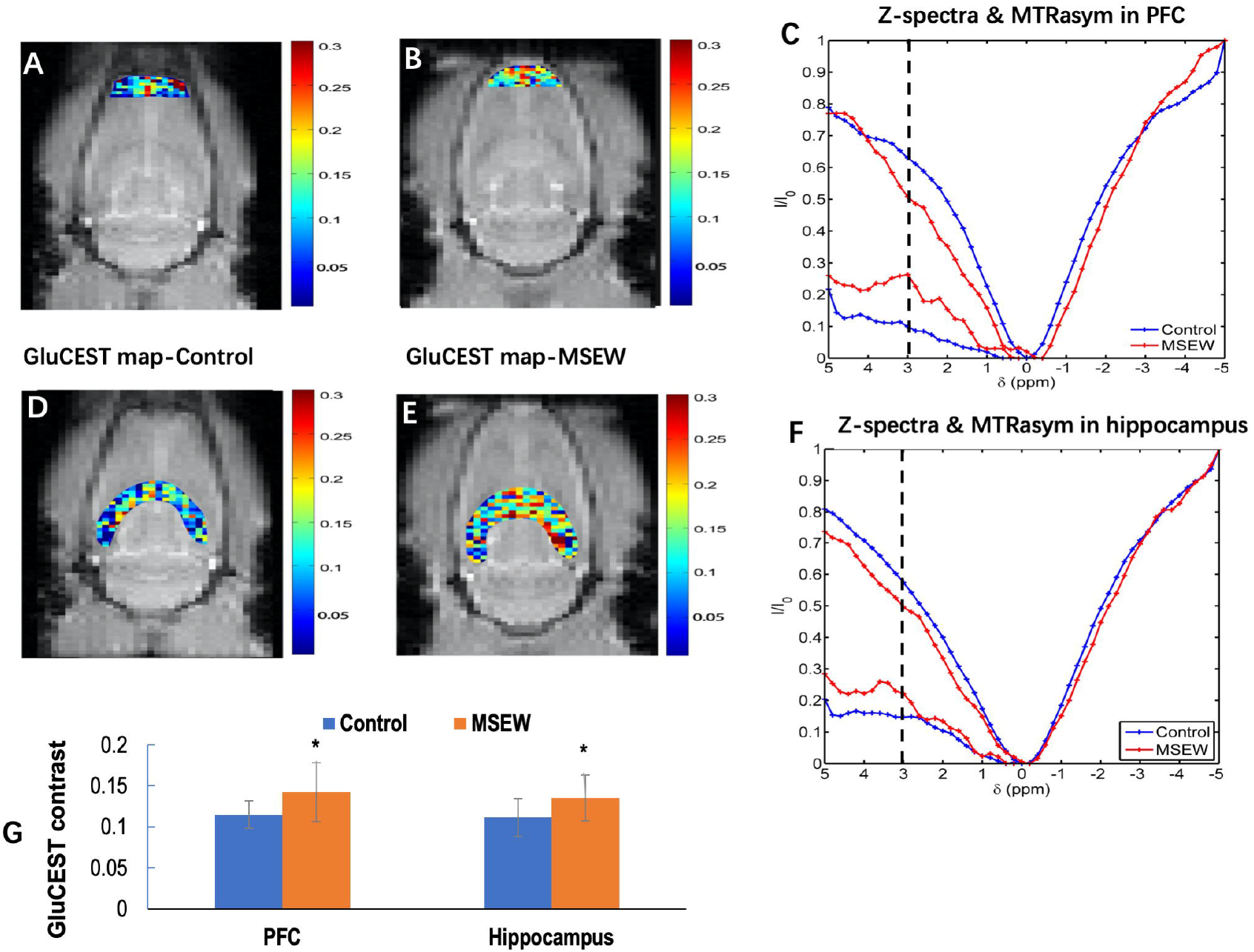
MSEW increased GluCEST contrast in PFC and dorsal hippocampus of rats. (**A**) & (**B**) The GluCEST map in PFC of Control and MSEW rats, respectively. (**C**) Corresponding z-spectra and MTRasym curves acquired at PFC of Control and MSEW rats. (**D**) & (**E**) The GluCEST map in dorsal hippocampus of Control and MSEW rats, respectively. (**F**) Corresponding z-spectra and MTRasym curves acquired at dorsal hippocampus of Control and MSEW rats. (**G**) Bar chart comparing the GluCEST contrasts in PFC and dorsal hippocampus of Control and MSEW rats. Data are expressed as mean ± SD. N = 13-14/group in two batches. Comparisons were made between Control and MSEW groups. *p<0.05.

### MSEW impairs the integrity of neuron-astrocyte and axon-myelin entities in rat brain: detected by ^1^H-MRS and DTI

Increased Glu levels in PFC and dorsal hippocampus of MSEW rats suggest a possibility that the neuron-astrocyte entity was damaged by the MSEW paradigm. To confirm this suggestion, we performed ^1^H-MRS scanning with rats following GluCEST scanning. The two volumes of interest (VOIs) were localized on the right side as shown in Fig. 3 (the inserts). Two representative ^1^H-MRS spectra obtained at frontal cortex were overlapped as shown in Fig. 3A (blue-Control; red-MSEW). The quantitative data of brain metabolites in this VOI of the two groups are shown in Fig. 3B indicating significantly higher levels of Glu, Glx (Glu+Gln), and NAA, but lower level of GABA, compared to Control group. Two representative ^1^H-MRS spectra obtained at the dorsal hippocampus were overlapped as shown in Fig. 3C (blue-Control; red-MSEW). The quantitative data of brain metabolites in this VOI of the two groups are shown in Fig. 3D indicating significantly higher levels of Glu, NAA, NAAG, and Tau, compared to Control group.

**Fig. 3.**
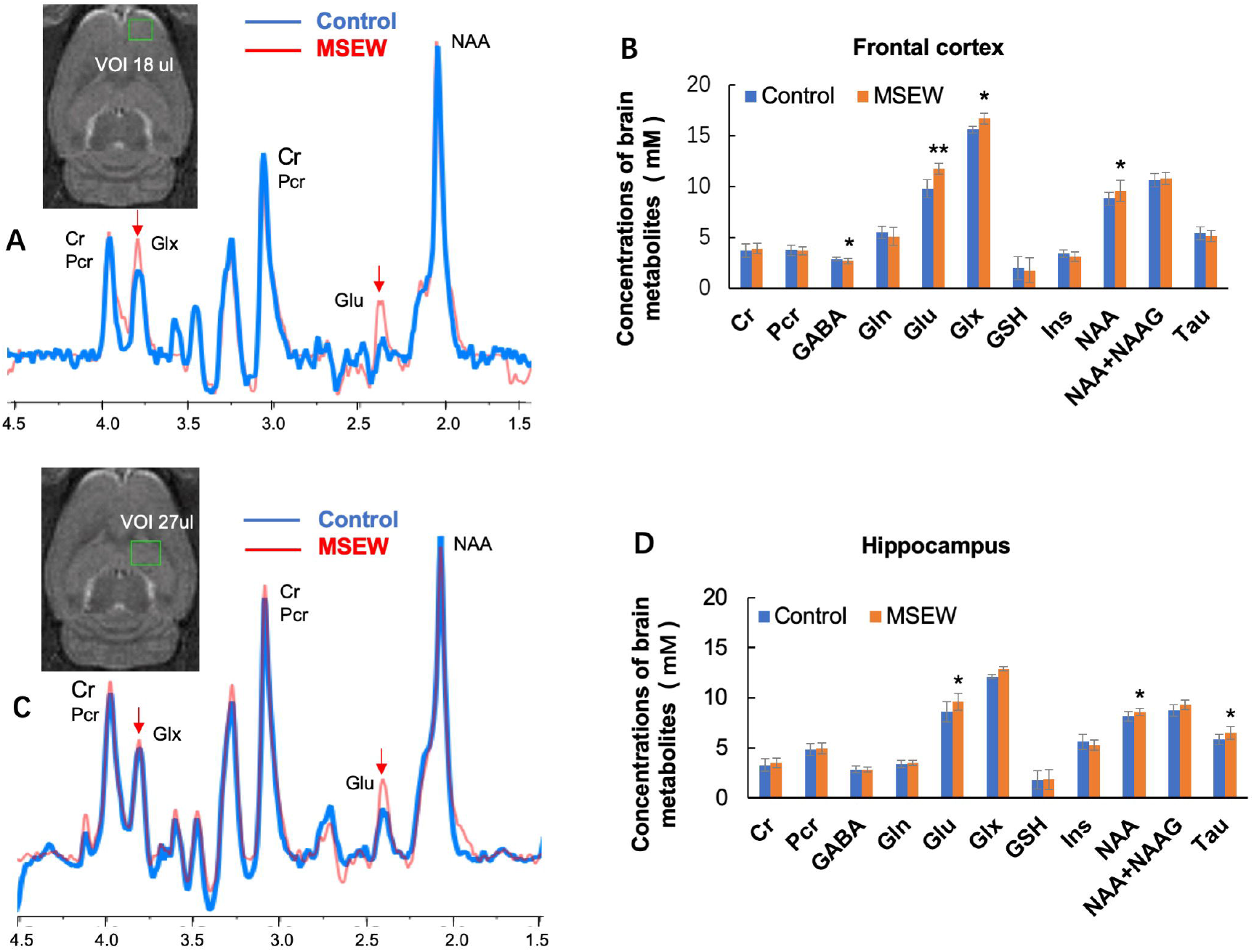
MSEW changed the neurochemical correlates of neuron-astrocyte integrity in PFC and dorsal hippocampus of the rat. (**A**) The overlapped two ^1^H-MRS spectra obtained at PFC of two rats in Control and MSEW groups, respectively (control: blue; MSEW: red). (**B**) Bar chart comparing levels of the brain metabolites in PFC of rats in the two groups. (**C**) The overlapped two ^1^H-MRS spectra obtained at dorsal hippocampus of two rats in Control and MSEW groups, respectively (control: blue; MSEW: red). (**D**) Bar chart comparing levels of the brain metabolites in dorsal hippocampus of rats in the two groups. Data are expressed as mean ± SD. N = 13-14/group in two batches. Comparisons were made between Control and MSEW groups. *p<0.05; **p<0.01.

In addition to NAA being used as the neurochemical correlate of axon-myelin entity as mentioned above, fractional anisotropy (FA) in DTI is commonly accepted as a biophysical index of myelin sheath integrity in the brain. It is a scalar value of the degree of directional diffusion within a voxel and linked to axon diameter, membrane permeability, and myelination. It is highly sensitive to microstructural changes within the location measured (Alexander et al., 2007). Therefore, we performed DTI with rats to look at if MSEW changed FA values in the brain. It was shown that FA values in frontal cortex, dorsal hippocampus, corpus callosum, and caudate putamen of MSEW rats were significantly lower than those in Control group (Fig. 4).

**Fig. 4.**
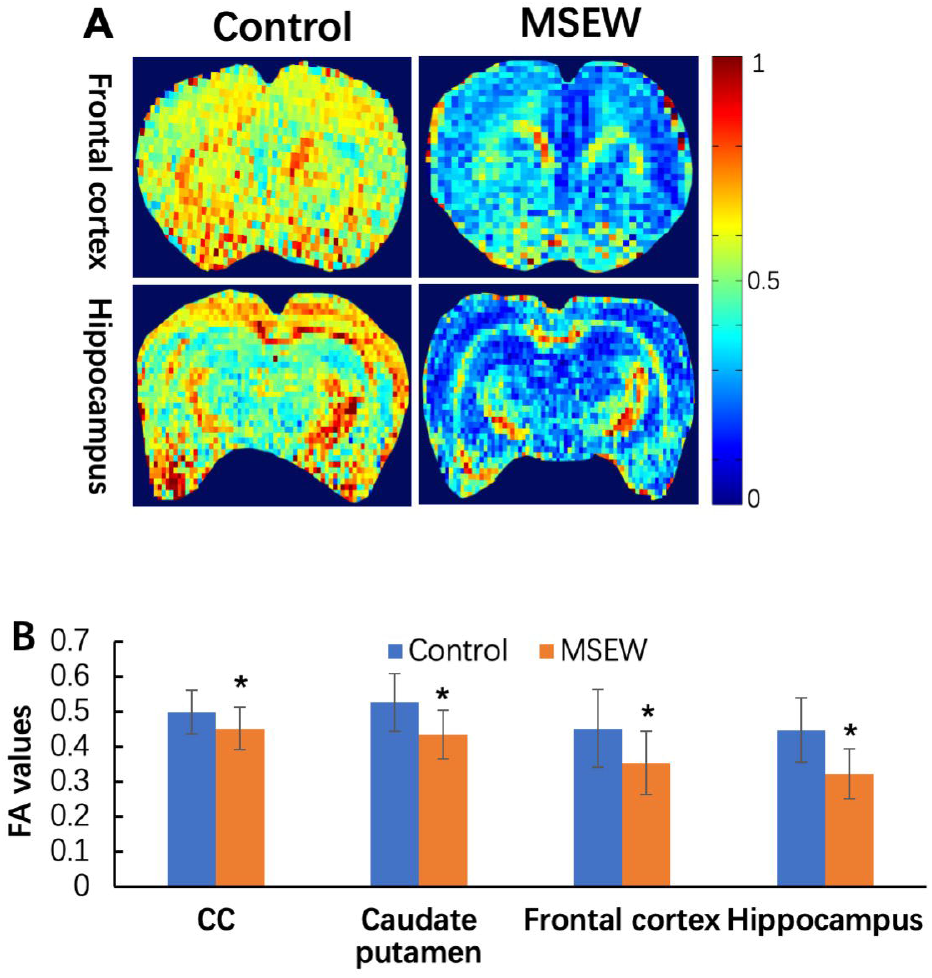
MSEW damaged the white matter integrity of rat brain. (**A**) Representative DTI maps from Control and MSEW rats showing MSEW-induced FA decreases in CC, caudate putamen, frontal cortex, and hippocampus. (**B**) Bar chart comparing FA values in all the measured brain regions of the two groups. Data are expressed as mean ± SD. N = 11-18/group in two batches. *p<0.05.

### MSEW disrupts glutamate-glutamine-GABA cycle in PFC of rats: the molecular underpinnings of impaired neuron-astrocyte integrity

To reveal the molecular underpinnings of impaired neuron-astrocyte entity shown by non-invasive neuroimaging methods, we measured the protein levels of ATP-α, GLT-1 (glial glutamate transporter 1), GLAST (the glutamate-aspartate transporter), PAG (phosphate-activated glutaminase), GAD65 (glutamate decarboxylase), and GS (glutamine synthetase) in the tissue sample of PFC. All these proteins involve in maintaining glutamatergic and GABAergic neurotransmission at optimal levels as reviewed in literature (Bak et al., 2006; Schousboe and Waagepetersen, 2005; Walls et al., 2015). Western blot analysis results showed lower levels of ATP-α and GLT-1, but higher levels of GAD65 and GS in PFC of MSEW rats compared to Control group; while no changes were found in levels of GLAST and PAG (Fig. 5A & 5B).

**Fig. 5.**
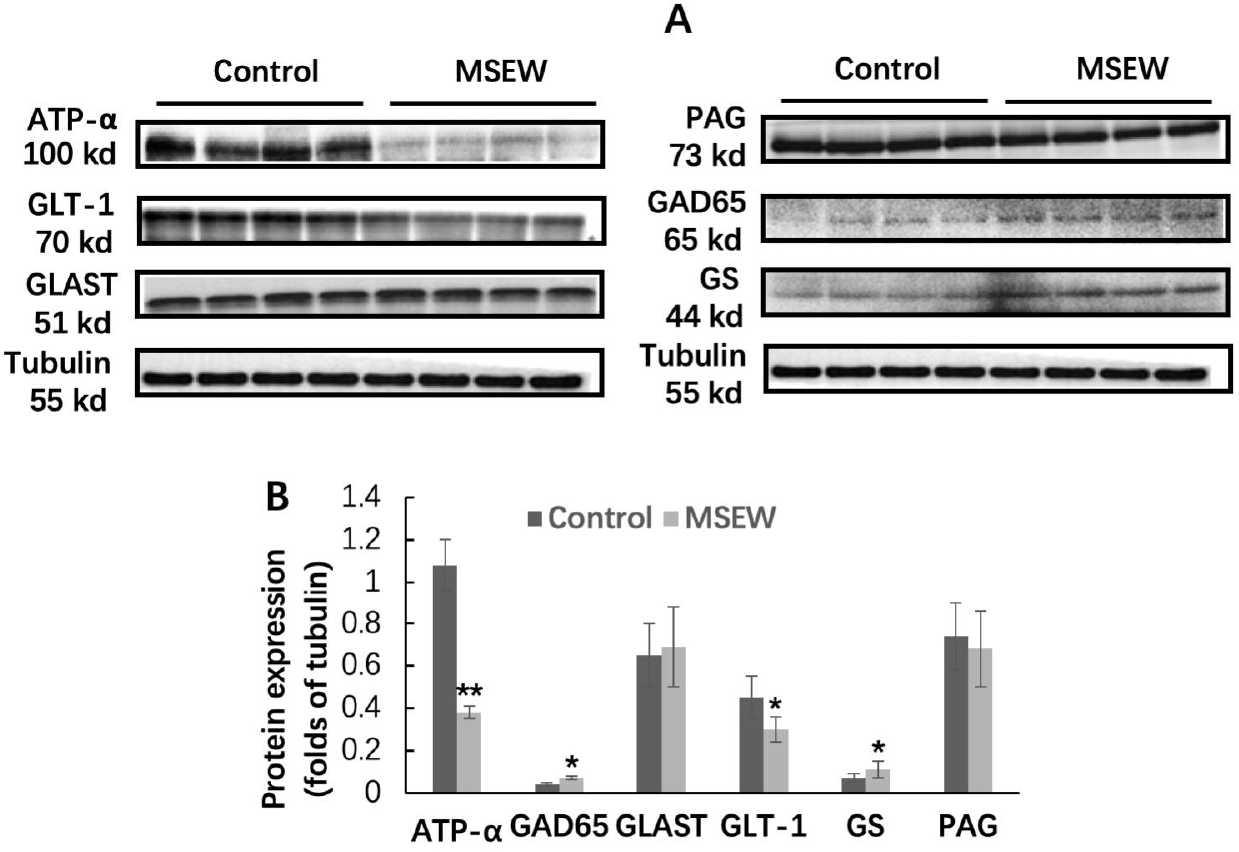
MSEW disrupted the glial metabolism of Glu in astrocytes. (**A**) Representative western-blot images of the target proteins (including ATP-α, GLT-1, GLAST, PAG, GAD65, and GS) in PFC of rats, along with that of **α**-tubulin as the reference. (**B**) Bar chart comparing levels of all the measured proteins of the two groups. Data are expressed as mean ± SD. N = 8/group. Comparisons were made between Control and MSEW groups. *p<0.05; **p<0.01.

### MSEW inhibits OLs maturation and myelination in PFC of rats: the cellular and molecular underpinnings of impaired neuron-oligodendrocyte integrity

To explore the cellular and molecular underpinnings of the MSEW-induced NAA increase and FA decrease in rat brain, we measured the protein levels of N-acetyltransferase 8-like (NAT8L) and aspartoacylase (ASPA) in tissue sample of PFC. NAT8L catalyzes the synthesis of NAA from acetyl-CoA and aspartate in neurons whereas ASPA works for the hydrolyzation of NAA into aspartate and acetate in OLs (Francis et al., 2016). Western-blot analysis showed lower levels of ASPA and NAT8L in PFC of MSEW rats compared to the Control group (Fig. 6), suggesting that the MSEW experience led to deficits in the expression of these two enzymes in OLs and neurons, respectively.

**Fig. 6.**
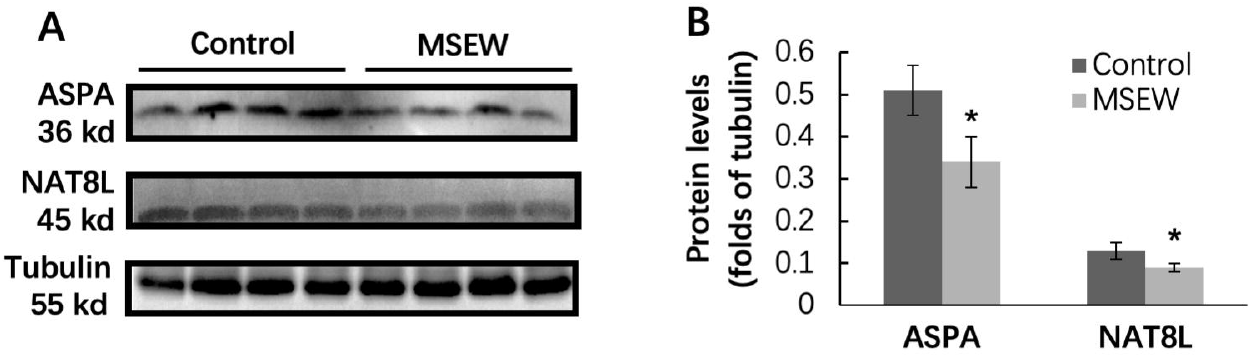
MSEW decreased levels of ASPA and NAT8L in PFC of the rat. (**A**) Representative western-blot images of ASPA and NAT8L in PFC of rats, along with that of **α**-tubulin as reference. (**B**) The bar chart comparing levels of ASPA and NAT8L in PFC of rats in Control and MSEW groups. Data are expressed as mean ± SD. N = 8/group. Comparisons were made between Control and MSEW groups. *p<0.05.

Concurrent NAA increase and ASPA decrease in PFC of MSEW rats suggest that the MSEW-induced functional deficit in OLs may happen in mature OLs or at advanced stages during OL development as ASPA activity increased markedly after PD7 and coincided with the time course for the onset of myelination in the rat brain (Bhakoo et al., 2001). To demonstrate this suggestion, we evaluated the mRNA expression of the transcription factors Olig2 and MRF (myelin-gene regulatory factor), as well as those of the myelin genes *Mbp* and *Plp1* in PFC of rats. Olig2 is an OL-lineage determination factor necessary for the OL lineage specification within the CNS (Lu et al., 2002; Zhou et al., 2002); whereas MRF is necessary for the generation of mature, myelin gene expressing OLs in the brain (Emery et al., 2009). RT-PCR analysis showed lower mRNA levels of *MRF, Mbp,* and *Plp1* in PFC of MSEW rats compared to Control group, but no difference was found in mRNA level of *Olig2* between the two groups (Fig. 7A). Western-blot analysis showed lower levels of MBP protein (both MBP17 and MBP21 isoforms) in PFC of MSEW rats compared to Control group, but no difference was found in PLP1 levels between the two groups (Fig. 7B & 7C). Moreover, we performed immunohistochemical staining using the antibody to MBP. The MBP immunoreactivity in the sub-regions of cingulate cortex (Cg1), prelimbic cortex (PrL), lateral orbital cortex (LO), and ventral orbital cortex (VO) of PFC was measured. The results indicate that MSEW rats had lower MBP immunoreactivity in PrL and VO compared to those in Control group, but that in Cg1 and LO was comparable between the two groups (Fig. 7D-F).

**Fig. 7.**
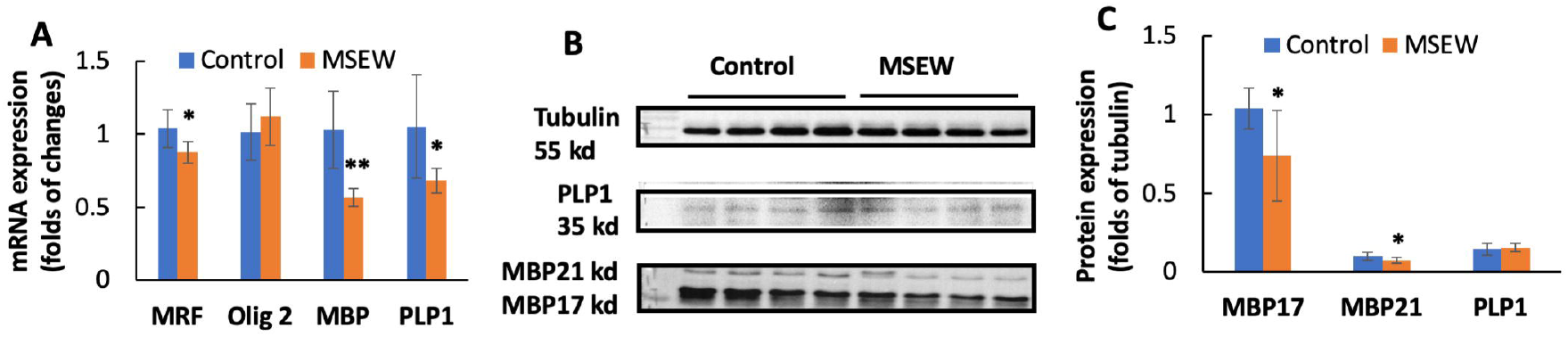
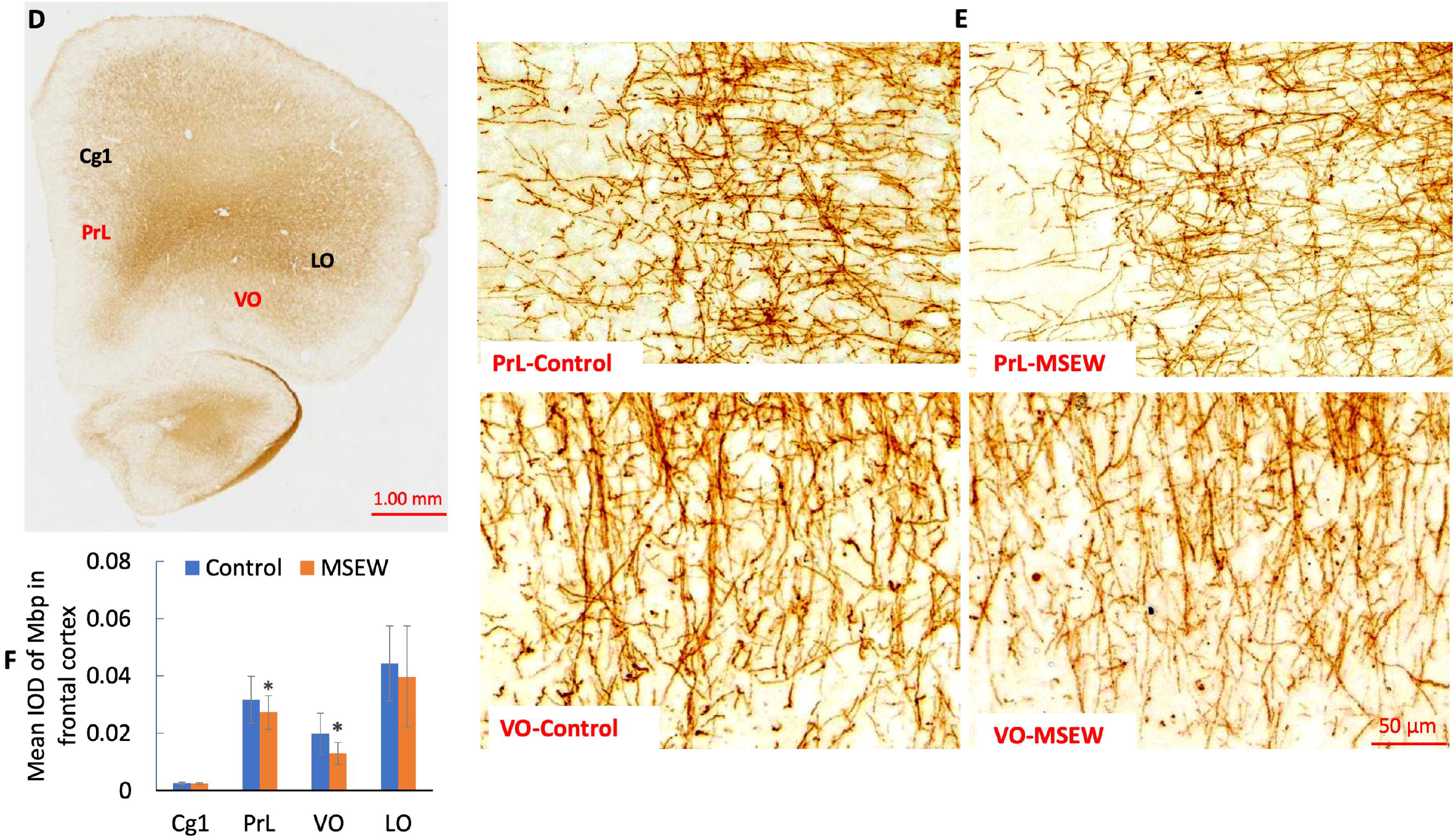
MSEW inhibited OLs maturation and myelination in PFC of rat brain. (**A**) The summarized data showing that MSEW decreases mRNA levels of *MRF, MBP, and PLP1* genes in PFC of rats, but has no effect on that of *Olig2* gene. (**B**) Representative western-blot images showing the immunoreactive bands of MBP (including MBP17 and MBP21 isoforms) and PLP1 proteins in PFC of rats. (**C**) Bar chart comparing levels of MBP (including MBP17 and MBP21 isoforms) and PLP1 proteins in PFC of Control and MSEW groups. (**D**) A representative immunochemical staining microphotograph taken at a lower magnification showing the subregions of Cg1, LO, PrL, and VO of PFC. (**E**) Representative microphotographs taken at higher magnifications showing the immunohistochemical staining with the antibody to MBP in PrL and VO in PFC of rats. (**F**) Bar chart comparing MBP immunoreactivity in the sub-regions Cg1, LO, PrL, and VO in PFC of Control and MSEW rats. Data are expressed as mean ± SD. N = 6-8/group. Comparisons were made between Control and MSEW groups. *p<0.05, **p<0.01. Abbreviations: Cg1, cingulate cortex; LO, lateral orbital cortex; PrL, prelimbic cortex; VO, ventral orbital cortex.

## Discussion

MSEW rats showed higher levels of anxiety and exploration in the open field test compared to those in the Control group. This is in line with a previous study showing that MSEW mice moved faster and thus covered more area than controls. The MSEW mice also spent less time in the center of the open field relative to controls (George et al., 2010). Increased anxiety-like behavior and activity levels were also seen in MSEW adult male mice of a recent study (Murthy et al., 2019). However, postnatal maternal separation (MS) demonstrated no clear and consistent effects on behavioral phenotypes in a variety of strains of mice in the other studies (Millstein and Holmes, 2007; Tan et al., 2017). Taken together, the results of present study and previous ones demonstrate that the MSEW mice and rats can be used as a robust model to mimic early life neglect and abuse in humans.

The GluCEST maps and regional analysis of GluCEST exhibited increased GluCEST contrast in PFC and dorsal hippocampus of MSEW rats relative to Control group, suggesting increased Glu concentrations in the two brain regions. In support of this suggestion, a previous study reported a significant increase in GluCEST contrast in rats 24 h after administration of modafinil, a drug known to increase Glu levels in the brain (Haris et al., 2014). In the other studies, restraint and swimming stress increased extracellular levels of endogenous excitatory amino acids, aspartate and Glu (Lowy et al., 1993; Moghaddam, 1993), and the stress-induced release of Glu is more pronounced in PFC of the rat as compared to other regions tested (Gilad et al., 1990; Moghaddam, 1993). All these point to a conclusion that GluCEST is a reliable non-invasive imaging method able to measure brain Glu levels in living subjects and has specific relevance to clinical research on patients with mental disorders thereof.

In addition to confirming Glu increase in PFC and dorsal hippocampus of MSEW rats, ^1^H-MRS revealed higher levels of NAA, but lower GABA in PFC relative to Control group. In the dorsal hippocampus, MSEW rats showed significantly higher levels of Glu, NAA, and Tau, but no change in GABA level compared to Control group. Increased Glu levels and decreased GABA in MSEW rats suggest a disrupted Glu-Gln-GABA cycle in the neuron-astrocyte entity in these rats (Xu et al., 2016). Also, these data are in line with the notion that Glu and GABA in PFC and hippocampus can be viewed as a general vulnerability to mood disorders like major depressive disorder and bipolar disorder (Lener et al., 2016). Adverse events during childhood increased risk of these mood disorders (Lupien et al., 2009). The pathologically elevated NAA in MSEW rats is indicative of impaired axon-myelin/neuron-OL entity as NAA level in the brain depends on NAA synthase in neurons and aspartoacylase in OLs. Higher levels of NAA indicate the degradation of NAA into L-aspartate and acetate was inhibited in OLs thus leading to deficit in acetate, a building block required for myelin lipid synthesis in OLs (Maier et al., 2015; Xu et al., 2016). Together, these data suggest that MSEW experience disrupted neuron-glia integrity in frontal cortex and dorsal hippocampus of rats.

Furthermore, we revealed the molecular changes underlying the MSEW-induced damage to neuron-astrocyte entity. First, MSEW paradigm induced concurrent decreases in levels of GLT-1 and ATP-α in PFC of rats. These two proteins were reported to co-localize within the final processes of astrocytes in two previous studies (Cholet et al., 2002; Roberts et al., 2014) and their interactions were demonstrated in the other studies (Bauer et al., 2012; Genda et al., 2011; Matos et al., 2013; Rose et al., 2009). Therefore, concurrent decreases in levels of GLT-1 and ATP-α must impair the uptake of Glu and the transport of it into astrocytes thus leading to Glu increase in the synaptic cleft and an initial decrease of Glu in astrocytes. Under this circumstance, the astrocytes in MSEW rats must increase *de novo* synthesis of Glu thus leading to a subsequent increase of Glu as revealed by GluCEST and ^1^H-MRS. Second, the conversion of Glu to Gln in astrocytes was sped up as evidenced by the upregulation of GS in the MSEW rats. Third, the protein levels of PAG, the neuron-specific enzyme responsible for the conversion of Gln to Glu, were comparable between the Control and MSEW groups, indicating that the conversion of Gln into Glu in glutamatergic neurons occurred normally, which may at least partly account for the comparable levels of Gln in MSEW and Control groups. Forth, MSEW rats showed higher level of GAD65, the enzyme catalyzing the conversion of Glu into GABA in GABAergic neurons. However, GABA level in PFC of MSEW rats was lower than Control group, suggesting that GAD65 increase in GABAergic neurons was not competent enough to maintain GABA at normal levels. Taken together, all these data demonstrate that MSEW paradigm in this study disrupted the glial metabolism of Glu in astrocytes and impaired neuron-astrocyte entity integrity thereof.

Concurrent elevated NAA levels and lower FA values in PFC and hippocampus of MSEW rats suggest that higher NAA level is a neurochemical index of axon-myelin entity integrity impairment in the brain as FA alterations are correlated to changes in myelination and axonal membrane integrity (Zatorre et al., 2012). This suggestion is consistent with previous studies showing: 1) MSEW significantly lowered FA values in medial septum and anterior commissure of mice (Carlyle et al., 2012); and 2) pubertal verbal stress in humans is associated with adult decreased FA values in several brain regions (Makinodan et al., 2012). In a more recent study, social defeat experience lead to concurrent NAA increase and MBP immunoreactivity decrease, but not OL loss, in medial prefrontal cortex (mPFC) of Balb/c mice (Zhang et al., 2016). All these previous studies and the present study demonstrate that NAA increase is indicative of myelination deficit in the brain. In support of this interpretation, NAA increment is considered to reflect myelin deficit in Canavan’s disease (Cakmakci et al., 2010; Wittsack et al., 1996), a recessive inherited disease characterized by NAA accumulation and myelin defect due to the deficiency of ASPA (Hoshino and Kubota, 2014). Like in the case of Canavan’s disease, MSEW rats in the present study showed decreased ASPA levels in PFC, suggesting functional deficit of OLs. Moreover, MSEW decreased NAT8L levels in PFC, suggesting the synthesis of NAA from acetyl-CoA and aspartate in neurons was also impaired. Via these actions, MSEW led to hypomyelination in the rat brain as evidenced by lower levels of MBP protein in PFC detected by means of Western-blot and immunohistochemical staining. Of note, MBP immunoreactivity in PrL and VO subregions of PFC was significantly lower in MSEW rats compared to Control group. These two subregions are in the lower medial and orbital prefrontal cortices in rodents and typically considered to be related to human and monkey Brodmann area 25, a component of ventromedial prefrontal cortex (vmPFC), a major player in social and affective functions (Hiser and Koenigs, 2018; Wise, 2008). Moreover, MSEW rats showed significantly lower levels of MRF mRNA, but no change in Olig2 mRNA levels, suggesting that the MSEW procedure affected OL maturation, but had no effect on OL differentiation. All the aforementioned data has confirmed the validity of NAA being used as a neurochemical index of axon-myelin integrity and revealed the mechanism by which MSEW impairs the myelinating function of OLs thus damaging the axon-myelin integrity in rat brain.

In summary, MSEW disrupted neuron-glia integrity in PFC and dorsal hippocampus of rats as evaluated by non-invasive neuroimaging methods showing changes in levels of Glu, NAA, and GABA in the brain regions, as well as global decrease in FA value. Underlying the disrupted neuron-astrocyte integrity in MSEW rats, changes happened in levels of those proteins that involve in maintaining glutamatergic and GABAergic neurotransmission at optimal levels. In support of damaged axon-myelin integrity, MSEW lowered levels of ASPA and NAT8L, as well as MBP levels and immunoreactivity in PFC. Along with these changes, MSEW increased locomotor activity and anxiety levels of rats.

## Materials and methods

### Animals

Female S-D rats at gestational week 2 were purchased from the animal center of the Southern Medical University (Guangzhou, China) and housed in an air-conditioned room at Shantou University Medical College. The animals had free accesses to food and water in the room with controlled temperature in the range of 23 ± 1 °C and a 12:12 h light cycle. The delivery day was defined as PD0. An even number up to ten pups (males) of each litter and their dam were culled to proceed to the next MSEW procedure or be used as controls. All animal handling and use were carried out in accordance with the guidelines set up by the Animal Care and Use Committee of Shantou University Medical College and approved by the committee.

### The MSEW procedure

The maternal separation started on PD2, by removing a pup from his/her dam and placing the pup in a small paper box (10×9×9 cm) for 4 hrs during PDs 2-5, and 6 hrs during PDs 6-16. During the separation periods, which started at the same time (8:00 am) every day, pups in cartons (one pup per carton) were kept at an infant incubator (YP-100; Ningbo David Medical Device Co., Ltd., Ningbo, China) which was kept well ventilated at a controlled temperature (34 °C during PDs 2-5, 32 °C during PDs6-9, 30 °C during PDs 10-14, and 28 °C during PDs 15-16) and humidity (60%). Before and after maternal separation, all pups in the MSEW group were brought back to the cage where their dam was living. Weaning occurred on PD 17 when a home-made soft diet (powdered rodent chow in tap water) was provided to the isolated pups. Starting on PD 22, the MSEW rats of a same litter were housed in group (10 pups/cage). The pups in Control group were raised by their dams under the standard laboratory condition as described before and weaned on PD 22. The body weight of all pups was weighed on PDs 7, 14, 21, and PD 30, respectively. During PDs 60-62, rats in the MSEW and Control groups were subjected to the open-field test.

### Open field test

Open field test was performed to measure the locomotor activity, exploratory behavior, and anxiety-related behaviors of rats. The wooden open-field box (100×100× 60 cm) was painted in black and sheltered by a blue drape in the behavioral test room, which was lighted with three fluorescents of 15 lumens. The experimental method has been described in a previous study (Shao et al., 2015). Briefly, each individual rat was placed in the center of the open-field box and allowed to move freely for 12 min, of which the first two minutes were defined to be the adaptation period and the data of this period was not included for analysis. A video tracking system (EthoVision XT 9.0; Noldus Information Technology, Wageningen, Netherlands) was used to monitor the tested rat. For each test, the travel distances on the whole arena and its central zone (the central part of 50 x50 cm), the travel speed, and time spent on the peripheral and central zones were recorded. The floor and inner walls of the box were cleaned with 70% ethanol after each test.

### Phantom scanning of MRI

All MRI scanning with phantom and animal were performed using an animal MRI scanner (7T, Agilent Technologies, Santa Clara, CA, USA) along with a 63 mm internal diameter standard proton transmission and reception volume coil. During MR imaging and spectroscopy measurements, animals were kept under anesthesia (1.5% isoflurane in 1 liters/min oxygen) and their body temperature maintained at 36–37°C.

A phantom was prepared as described in a previous study to optimize CEST MRI procedure (Zhuang et al., 2019). Briefly, 1% agarose was added into a phosphate-buffered saline (PBS) solution in test tubes (5 mm in diameter) and heated by a microwave. The mixture was then immersed in a water bath at 45 °C. Glu (Sigma-Aldrich, St Louis, MO, USA) was added at the various concentrations of 0, 3, 6, 9, 12, and 15 mM, and the pH was titrated to 7.0. The tubes were inserted into a phantom holder filled with 1% agarose gel to minimize susceptibility inhomogeneity. In addition, another phantom consisting of tubes with solutions of different metabolites at their physiological concentrations (Asp, 2 mM; Cr, 6 mM; GABA, 2 mM; Gln, 2 mM; Glu, 10 mM; NAA, 10 mM). All these chemicals were purchased from Sigma-Aldrich (St Louis, MO, USA) and dissolved in a 1% agarose gel at the pH of 7.0.

### Diffusion tensor imaging

Diffusion tensor imaging was performed for anesthetized individual rat placed in the prone position with the head fixed on a palate holder equipped with an adjustable nose cone. Respiration and body temperature were monitored using an MRI compatible small animal monitor system (SA Instruments, Inc., Stony Brook, NY, USA). The body temperature of rat was maintained at 37°C using a water-heated animal blanket. High resolution anatomical T_2_-weighted images were acquired with multi-slices multi-echoes (MSME) sequence (TE/TR = 40/3000 ms, average =1, repetition =1, matrix = 256 × 256, slice thickness = 2.0 mm) in coronal (10 slices), sagittal (8 slices) slices. 4 sub-regions in 2 sagittal slices were orientated and used for accurate delineation of the target regions. DTI scanning was carried out by using a fast spin echo (FSE) diffusion-weighted sequence with the parameters of: TR = 2000 ms, repetitions = 1, average = 8, data matrix = 192 × 128, FOV = 45 × 45 mm, slice = 8, thickness = 1 mm, gap = 0.2 mm, b = 1,000 s/mm^2^. DTI data were processed using Diffusion Toolkit in Track Visual software according to a previous study^18^. The axes of the ellipsoid are described by a set of eigenvectors and eigenvalues (λi) to calculate FA values obtained in four ROIs including CC, caudate putamen, frontal cortex, and hippocampus in corresponding target slices. All these DTI measures were obtained and analyzed by AFNI (Analysis of Functional NeuroImages) software.

### GluCEST imaging

GluCEST imaging was done as described above in DTI subsection. T_2_-weight images were acquired. The second one of the horizontal slices passing through the frontal cortex and dorsal hippocampus was selected for GluCEST image. A custom-programmed segmented RF spoiled gradient echo (GRE) pulse sequence was used. The sequence parameters were as follows: FOV = 50 mm × 50 mm. TR = 17.58 ms; TE = 2.77 ms, flip angle = 15 deg, average = 8, matrix = 128 x 64, slice thickness = 2 mm. CEST images were collected using a 1 s saturation pulse at B1 of 250 Hz (5.9 μT) at multiple frequencies in the range - 5 to + 5 ppm, with a step size of 0.2 ppm. The scan time was 7 m 57 s for each animal. Control images (S_0_) were obtained at an offset of 300 ppm for normalization, and GluCEST was measured at 3 ppm.

For MTR mapping, the images at 20 ppm and 100 ppm were collected on the same brain slice with a saturation power of 250 Hz and a saturation duration of 1 s. An image at 100 ppm was considered as the magnetization off image.

All GluCEST image processing and data analysis were performed using programs employed in previous studies (Haris et al., 2013). CEST images were corrected by B0 and B1 map and z-spectra were obtained from the normalized images for the ROI outlined. CEST contrast was calculated according to the Equation:

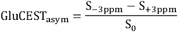

Where S_±3ppm_ was the magnetization obtained with saturation at a ‘+’ or ‘-’ 3ppm offset from the water resonance, in which +3ppm was consider as frequency offset of Glu for chemical exchange and saturation transfer. S_o_ is the magnetization with a saturation pulse applied at 100 ppm, as a control image for data normalization.

MTR maps were also computed using Equation:

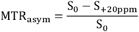

Where S_+20Ppm_ was the magnetization of macromolecules acquired with saturation pulse at +20 ppm.

### Proton MRS

Proton MRS was done as described in GluCEST subsection. A single voxel spectrum was acquired with point resolved spectroscopy (PRESS) using a vendor (Varian) provided pulse sequence. The parameters were as follows: TR = 3000 ms, TE1 = 7.61 ms, TE2 = 6.01 ms, TE total = 13.62 ms, average = 12, dummy scan = 2, real points = 4095, spectral width = 4005 HZ, slice thickness = 2 mm, voxel size in hippocampus = 3.0 x 3.0 x 3.0 mm and in frontal cortex = 3.0 x 3.0 x 2.0 mm. The scan time was 21 min. 3D voxel shimming was performed to obtain localized water line width values of lower than 12 Hz. Water suppression was performed by the variable pulse power and optimized relaxation delays method (VAPOR). An unsuppressed water spectrum was also acquired using the same parameters for normalization. Both water-suppressed and water reference spectra were analyzed by LC-model. The metabolites were quantified under the condition of Cramér-Rao lower bound (CRLB) □ 10% in experiments except GABA □ 20%. MRS spectra was fitted by MestReNova software.

### RT-PCR analysis

Tissue samples were lysed using Trizol and chloroform. The extracted RNA was quantified by microplate reader (Tecan, Infinite^®^ M1000 Pro, China). Then, the RNA sample was diluted into 140 ng/μL and purified with the Primescript TM RT regent kit (Takara, RR047). cDNA was generated using SYBR^®^ Premix Ex Taq™ II kit (Takara, RR820A). Quantitative real-time PCR was run on the Applied Biosystems7500 Real Time PCR System (Applied Biosystems, Foster City, CA, USA) by following the 2 steps real time RT-PCR procedure and using primers listed in Table 1. Each CT value acquired from quantitative RT-PCR was calculated as average of triplicate samples from one rat (8 in each group). GADPH gene was used as the reference gene. Relative gene expression levels were analyzed using the 2^ (-ΔΔ CT) method.

### Western-blot analysis

The PFC samples of rats were homogenized, and proteins were extracted using a Tris–EDTA lysis buffer (1% Triton X-100, 10% glycerol, 20 mM Tris, pH 7.5, 1 mM EDTA) containing freshly added protease inhibitor cocktail (Sigma-Aldrich, St. Louis, MO, USA). After protein determination using a BCA kit, sodium dodecyl sulfatepolyacrylamide gel electrophoresis and Western blotting were performed. The antibody against β-tubulin (1:1000) (Sigma-Aldrich, St. Louis, MO, USA) was used as an internal control for the concentration of protein loaded, while the primary antibodies against ATP-α (1:1000), GLT-1 (1:5000), GLAST (1:2000), GAD65 (1:1000), GS (1:500), or PAG (1:1000) were used to detect the corresponding target proteins. After incubation in a secondary antibody solution (anti-rabbit antibody, 1:1000), the immunoreactive bands were visualized using an ECL detection kit (Amersham Biosciences, Buckinghamshire, UK). Image Lab software version 5.0 (Bio-Rad) was used for image acquisition and densitometric analysis of the immunoreactive bands. The data were expressed as folds of **α**-tubulin band in a corresponding lane.

### Immunohistochemical staining

Immunohistochemical staining was performed as described previously (Zhang et al., 2018). The brain sections (20 μm thickness) were blocked with 5% goat serum (GeneTex, Alton PkwyIrvine, CA) in 0.01 M PBS containing Triton X-100 (0.1%) for 1 h at room temperature, followed by the incubation with the primary antibody against MBP(1:200; Abcam, Cambridge, UK) overnight at 4°C. After rinsed three times with PBS, the sections were incubated with HRP-conjugated secondary antibody (goat antirabbit IgG, 1:1000; Beyotime Biotechnology, Shanghai, China) at 37 °C for 1.5 h at room temperature. After rinsed in PBST, the sections were visualized by adding diaminobenzidine solution as recommended by the manufacturer (Zhongshan Gold Bridge Biology Company, Beijing, China). After rinsed in PBS, the sections were mounted onto the glass slides, dehydrated in gradient ethyl alcohol and cleared in xylene. After mounted by neutral balsam, immunohistochemical staining was observed and recorded with a Zeiss microscope (Zeiss instruments Inc., Germany) and analyzed with the Image-Pro-Plus 6.0 software (Media Cybernetics, Rockville, Maryland). The negative control staining was done following the same procedure except for the deletion of the primary antibody. No positive MBP immunoreactive staining was seen on the negative control sections.

### Analysis of immunohistochemical staining

As described previously (Zhang et al., 2018), the quantitative analysis of immunohistochemical staining was performed with the sub-regions of Cg1, LO, PrL, and VO of PFC as ROIs. For each ROI, three sections were chosen. For each section, three images were recorded under the same conditions. The Image-Pro Plus 6.0 software was used to automatically read out the measurement values. The data of MBP-like immunoreactivity (IOD, integrated optical density) were expressed as percent of Control group.

### Statistical analysis

SPSS17.0 (IBM Corp., Armonk, NY, USA) was used to analyze all the data which were expressed as mean ± SD. The Shapiro–Wilk test was used for normality of data distribution. Independent student t-test was used to compare mean values of two groups. The significant threshold was set at 0.05.

## Acknowledgements

This work was supported by funds from Shantou University Medical College, Li Kashing Foundation (H.X.; grant no. 432095 02) and the Science and Technology Bureau of Guangdong Province, China (H.X.; grant no. 2016A030313067, Q.H. and H.X.; grant no. 2015KGJHZ018).

## Author contributions

H.Z., W.W., Q.H., and H.X. conceived the idea. H.Z., Z.S., Z.D., and X.C. performed DTI, GluCEST, and ^1^H-MRS scanning and analyzed the neuroimaging data. G. Y., X.Z., R.W., and H.X. instructed the neuroimaging experiments and interpreted the neuroimaging data. H.Z., Z.Y., and S.X. performed behavioral tests, immunohistochemical staining, RT-PCR, and Western-blot experiments. H.Z and H.X. interpreted and integrated all the data. Q.H., R.W., and H.X. supervised the research. H. Z. and H.X. wrote the manuscript with input from all coauthors. All authors approved the final version of the manuscript.

---

**Supplementary Table 1.**
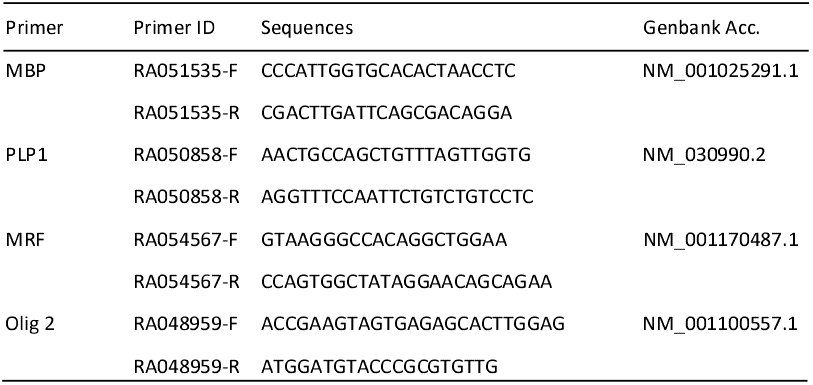
Primer sequences used for real-time RT-PCR

