## Supplementary figures and images for "Neuron-glia Integrity: Functional Assessment, Molecular Underpinnings, and Implication for Higher Brain Functions"

### Supplemental Fig. 1

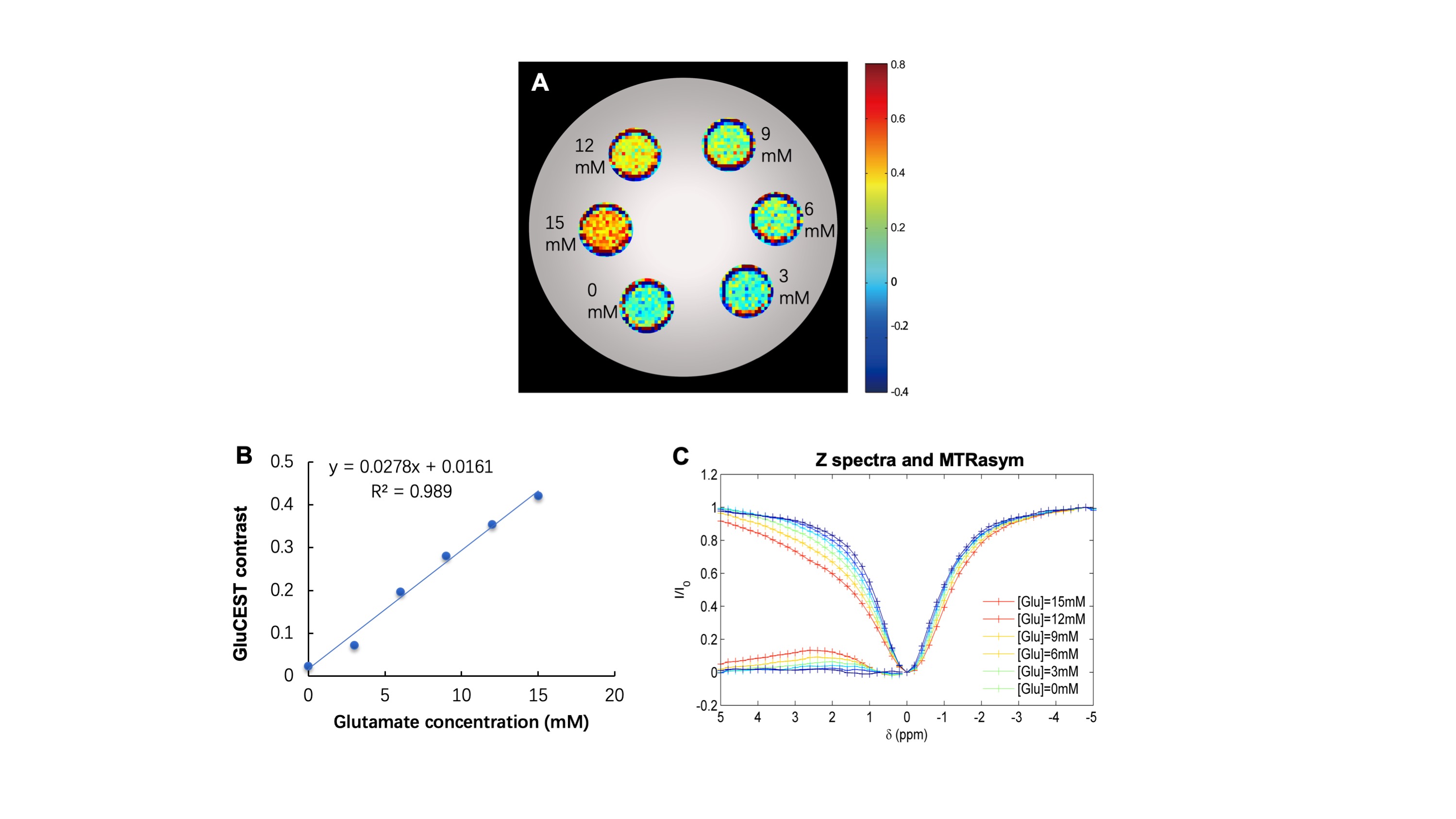

### Supplemental Fig. 2

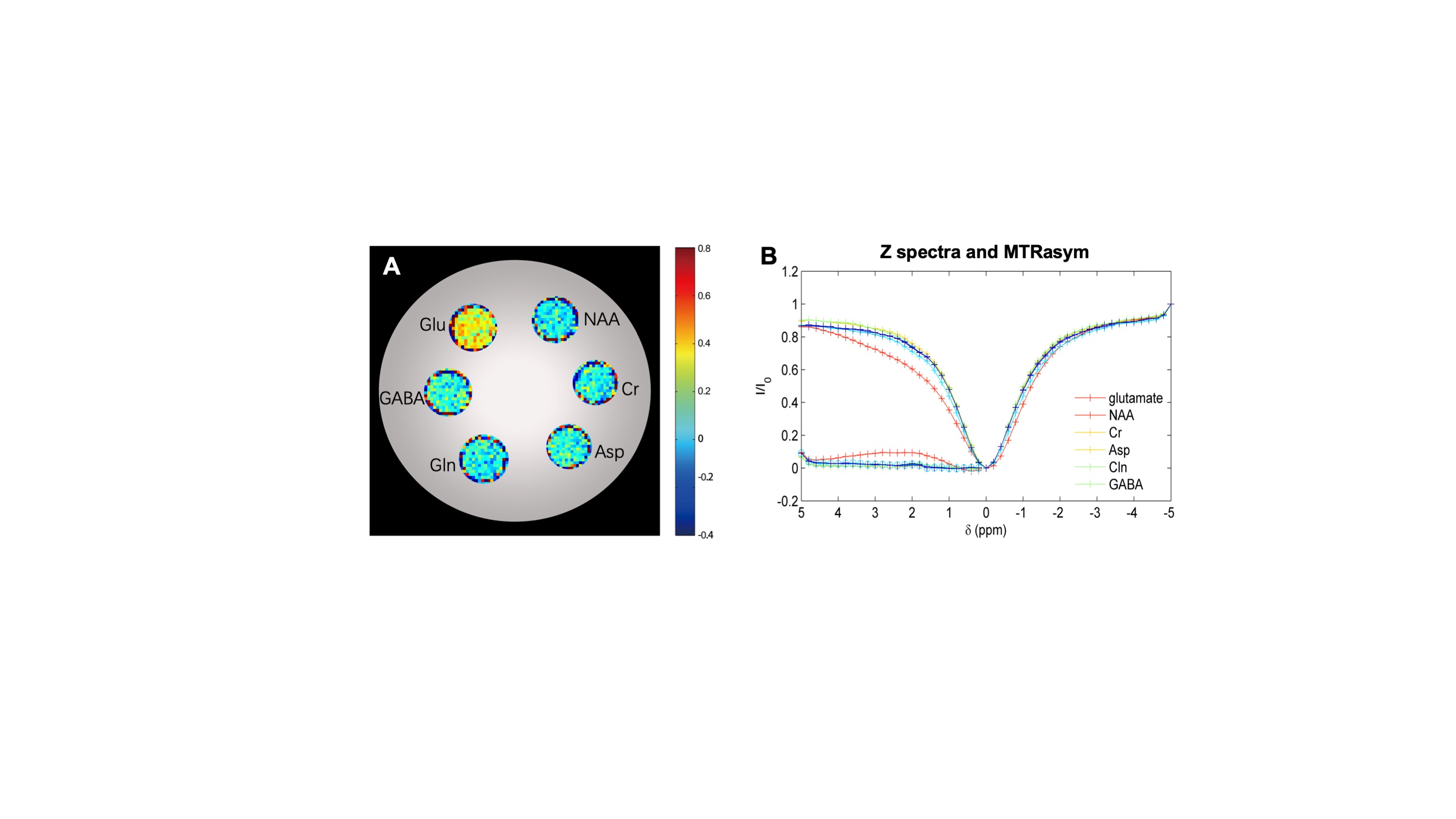

### Supplemental Fig. 3

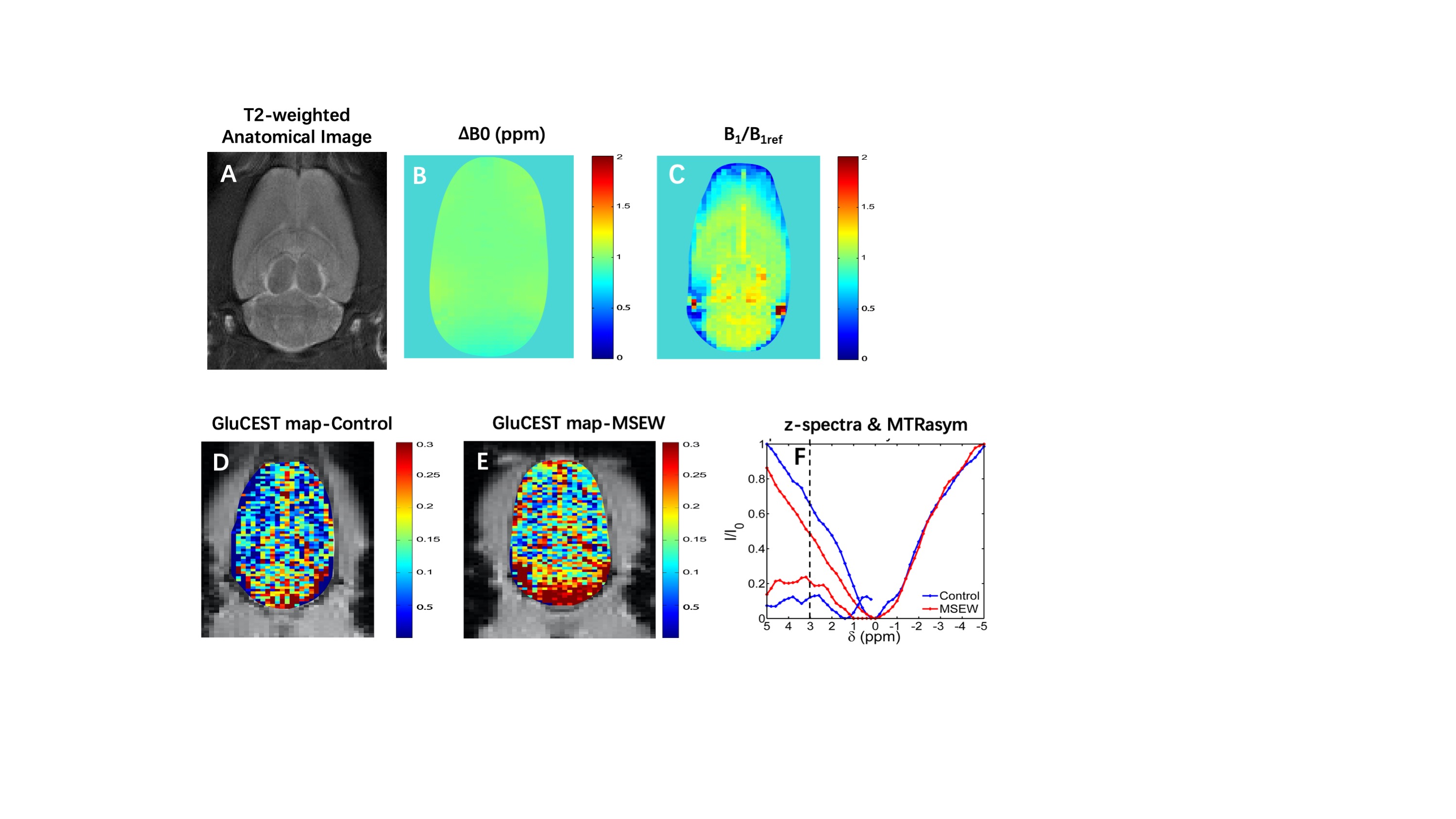
